# Biphasic unbinding of Zur from DNA for transcription (de)repression in Live Bacteria

**DOI:** 10.1101/434738

**Authors:** Won Jung, Peng Chen

## Abstract

Transcription regulator on-off binding to DNA constitutes a mechanistic paradigm in gene regulation, in which the repressors/activators bind to operator sites tightly while the corresponding non-repressors/non-activators do not. Another paradigm regards regulator unbinding from DNA to be a unimolecular process whose kinetics is independent of regulator concentration. Using single-molecule single-cell measurements, we find that the behaviors of the zinc-responsive uptake regulator Zur challenges these paradigms. Apo-Zur, a non-repressor and presumed non-DNA binder, can bind to chromosome tightly in live *E. coli* cells, likely at non-consensus sequence sites. Moreover, the unbinding from DNA of its apo-non-repressor and holo-repressor forms both show a biphasic, repressed-followed-by-facilitated kinetics with increasing cellular protein concentrations. The facilitated unbinding likely occurs via a ternary complex formation mechanism; the repressed unbinding is *first-of-its-kind* and likely results from protein oligomerization on chromosome, in which an inter-protein salt-bridge plays a key role. This biphasic unbinding could provide functional advantages in Zur's facile switching between repression and derepression.

## INTRODUCTION

Transcriptional regulation in cells is generally orchestrated by regulators, which, upon binding to operator sites, either block the binding of RNA polymerase (RNAP) leading to repression (i.e., repressors) or recruit RNAP leading to activation (i.e., activators)^1, 2^. One mechanistic paradigm for these regulators is an on-off model in which they bind to their cognate operator sites tightly, while their corresponding non-repressor/non-activator forms have insignificant affinity to DNA and stay predominantly in the cytoplasm. Some exceptions recently emerged. For example, IscR, a member of the MarA/SoxS/Rob family of transcription regulators in *E. coli*, is a repressor in its holo-form (i.e., containing a Fe-S cluster); its apo-form, generally thought to not bind DNA, was shown to bind DNA motifs different from its holo-repressor form^3, 4^.

Derepression or deactivation subsequently comes from the unbinding of the regulator from the operator site. Here another mechanistic paradigm exists regarding the kinetics of regulator unbinding, which is presumed to be a unimolecular reaction (i.e., spontaneous unbinding), whose first-order rate constant is independent of surrounding regulator concentration. However, recent *in vitro* single-molecule and bulk measurements uncovered facilitated unbinding, in which the first-order unbinding rate constant increases with increasing protein concentrations^5^. These proteins include nucleoid associated proteins that bind double-stranded DNA nonspecifically^6^, replication protein A that binds single-stranded DNA nonspecifically^7^, and DNA polymerases^8, 9^. We also discovered that CueR and ZntR, two MerR-family metal-sensing transcription regulators that bind to their cognate promoter sequences specifically, also show facilitated unbinding^10^. Using single-molecule tracking (SMT) and single cell quantification of protein concentration (SCQPC) that connect protein-DNA interaction kinetics with cellular protein concentrations quantitatively, we further showed that the facilitated unbinding of CueR and ZntR also operate in living *E. coli* cells^11^. A mechanistic consensus emerged, involving multivalent contacts between the protein and DNA^5^, which enables the formation of ternary complexes as intermediates that subsequently give rise to concentration-enhanced protein unbinding kinetics.

Here we report a SMT and SCQPC study of Zur, a Fur-family homodimeric zinc-uptake regulator, whose Zn^2+^-bound holo-form binds to its cognate operator site with nM affinity and represses the transcription of zinc uptake genes under zinc stress^12–15^; its apo-form is a non-repressor. We found that in living *E. coli* cells, Zur’s interactions with DNA challenge the above two paradigms. First, apo-Zur, long thought to not bind DNA, can bind to chromosome tightly, likely at non-consensus sites. Second and more strikingly, the unbinding of both apo- and holo-Zur from chromosome not only show facilitated unbinding with increasing cellular protein concentrations, but also exhibit repressed unbinding at lower concentrations, giving a first-of-its-kind biphasic unbinding behavior. The repressed unbinding of Zur likely stems from Zur oligomerization on DNA, where an inter-dimer salt bridge plays a key role, and it likely facilitates transcription switching between repression and depression in cells.

## RESULTS

### SMT and SCQPC identify a tight DNA-binding state for both holo- and apo-Zur in cells

To visualize individual Zur proteins in *E. coli* cells, we fused the photoconvertible fluorescent protein mEos3.2^16, 17^ to its C-terminus creating Zur^mE^, either at its chromosomal locus to have physiological expression or in an inducible plasmid in a Δ*zur* deletion strain to have a wider range of cellular protein concentrations (Methods). This Zur^mE^ fusion-protein is intact and as functional a repressor as the wild-type (WT) in the cell under Zn stress growth conditions (Supplementary Fig. 1a-b).

Using sparse photoconversion and time-lapse stroboscopic imaging, we tracked the motions of photoconverted Zur^mE^ proteins individually in single *E. coli* cells at tens of nanometer precision until their mEos3.2 tags photobleached (Fig. 1a). This SMT allows for measuring Zur^mE^’s mobility, which reports on whether the molecule is freely diffusing in the cell or bound to DNA. We repeated this photoconversion and SMT cycle 500 times for each cell, during which we counted the number of tracked protein molecules. We then used the SCQPC protocol to quantify the remaining number of Zur^mE^ protein molecules in the same cell^11^, eventually determining the Zur^mE^ concentration in each cell (i.e., [Zur^mE^]_cell_). This single-cell protein quantitation allowed for sorting the cells into groups of similar protein concentrations and subsequently examining protein-concentration-dependent processes, without being limited by the large cell-to-cell heterogeneity in protein expression.

**Fig. 1.**
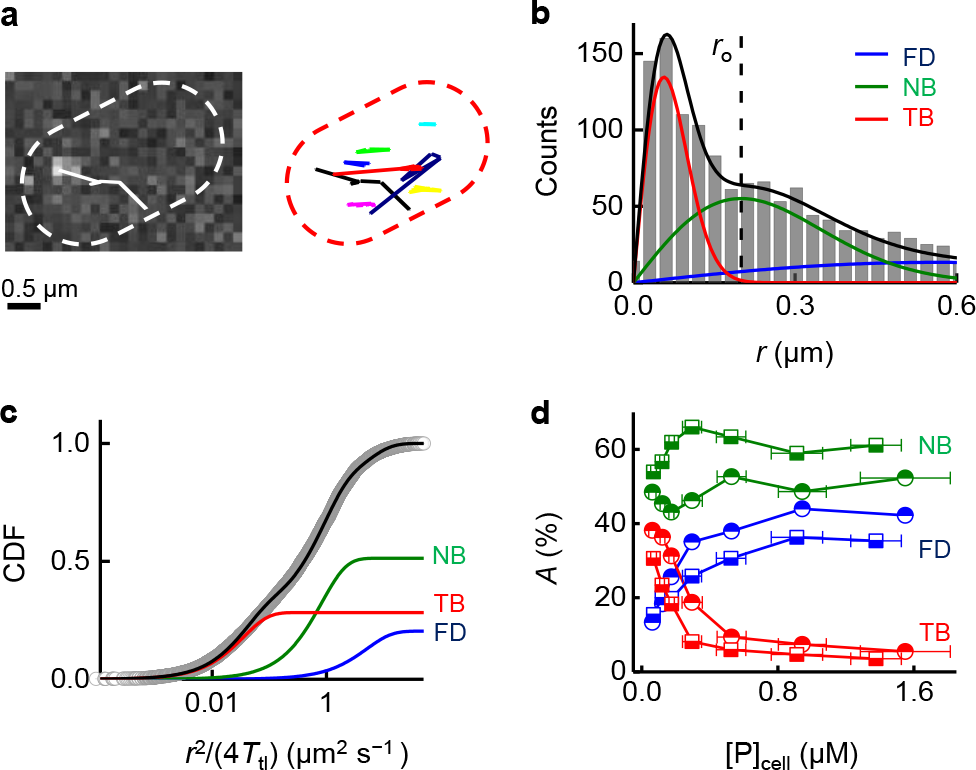
SMT of Zur in living cells. **a**, Left: exemplary fluorescence image of a single 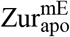 protein in a live *E. coli* cell overlaid with its position trajectory (solid line). Right: overlay of many trajectories. Dash lines: cell boundary. **b**, Histogram of displacement length *r* per time-lapse (40 ms) of > 1,400 tracked 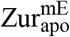 proteins at 124 ± 15 nM. Solid lines: the overall fitted distribution (black), and the resolved FD (blue), NB (green), and TB (red) diffusion states (Supplementary Note 4). Vertical dashed line: *r*o = 0.2 μm for extracting residence times as in Fig. 2a. **c**, Cumulative-distribution-function (CDF) of *r* (plotted against 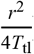) as in **b**. Lines: overall fit (Eq. (3)) and three resolved diffusion states with effective diffusion constants (and fractional populations): *D*_FD_ = 5.0 ± 0.5 μm2 s-1 (21.7 ± 0.4%), *D*_NB_ = 0.8 ± 0.05 μm^2^ s^−1^ (48.8 ± 0.4%), and *D*_TB_ = 0.040 ± 0.003 μm^2^ s^−1^ (30.1 ± 0.5%). **d**, Fractional populations of FD, NB, and TB states for 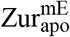 (half-solid squares) and 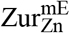 (half-solid circles) vs. the cellular protein concentrations.

We first examined 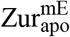 whose regulatory Zn-binding site was mutated (i.e., C88S) to make it permanent apo and a non-repressor^15^ (Supplementary Fig. 1b). To quantify its mobility in cells, we determined the distribution of its displacement length *r* between successive images and the corresponding cumulative distribution function (CDF) of *r* for each cell group having similar cellular 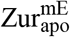 concentrations (Fig. 1b-c). Global analysis of these CDFs across all cellular protein concentrations resolved minimally three Brownian diffusion states with *effective* diffusion constants of ~5.0 ± 0.5, 0.82 ± 0.05, and 0.040 ± 0.003 μm^2^ s^−1^ (Fig. 1b-c; Methods). No subcellular localization or protein aggregation was observed; therefore, these two aspects are not the reasons for the presence of these three diffusion states. On the basis of their diffusion constants and previous studies of transcription regulator diffusion in *E. coli* cells^11^, ^18–21^, we assigned the fastest diffusion state as 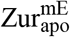 proteins freely diffusing (FD) in the cytoplasm, the medium diffusion state as those nonspecifically bound (NB) to and moving on chromosome, and the slowest state as those tightly bound (TB) to the chromosome, whose small effective diffusion constant (~0.040 μm^2^ s^−1^) reflects chromosome dynamics^19, 22^ and measurement uncertainties. Control measurements on the free mEos3.2 further support the assignment of the FD state, as we reported^11^.

The resolution of CDFs of *r* also gave the fractional populations of the three states across the range of cellular protein concentrations (Fig. 1d). With increasing 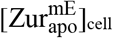, the fractional population of the FD state increases, while that of the TB state decreases. These trends further support their assignments because, with increasing cellular protein concentrations, more proteins compete for the limited number of tight binding sites on chromosome, leading to smaller fractional populations of the TB state and larger fractions of the FD state.

The presence of a significant fraction of the tight DNA-binding state, even at low cellular protein concentrations, is surprising for 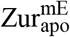 (e.g., ~32% at 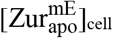 ~ 60 nM; 1 nM in an *E. coli* cell corresponds to ~1 protein copy), as apo-Zur is a non-repressor. Furthermore, previous gel shift assay showed that *E. coli* apo-Zur does not bind to operator sites (i.e., *K*_D_ > 300 nM at the *znuABC* promoter)^15^, and for *B. subtilis*, its apo-Zur’s binding affinity to operator sites is ~1000 times weaker than its holo-form^23^. We hypothesized that the TB state of 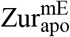 likely comes from its binding to non-operator sites (i.e., non-consensus sequence sites; see later).

We next examined Zur^mE^ in cells stressed with 20 μM Zn^2+^ in the medium. This Zn^2+^ concentration can evoke maximal repression of *zur* regulons (Supplementary Note 2.3). Therefore, most of Zur proteins in the cell should be metallated, mimicking the holo repressor form (i.e., 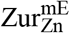). The same three diffusion states are resolved in the CDFs of *r* across all cellular protein concentrations (Supplementary Note 4.2). In contrast to the case for 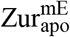, the TB state of 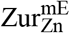 is expected here because holo-Zur binds specifically to consensus operator sites within Zur-regulated promoters. Expectedly, the fractional population of the FD state of 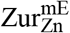 increases with increasing 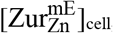, whereas that of the TB state decreases (Fig. 1d).

### Concentration-dependent biphasic unbinding kinetics of Zur from DNA

To probe Zur-DNA interaction dynamics, we examined the *r* versus time *t* trajectories of individual Zur proteins inside cells. These trajectories show clear transitions between large and small *r* values (Fig. 2a): the small *r* values are expected to be dominated by instances of Zur tightly bound to chromosome (i.e., TB state). We set an upper threshold *r*_0_ (= 0.2 μm), below which >99.5% of the TB states are included based on the resolved distributions of *r* (Fig. 1b), to select these small displacements and obtain estimates of the individual residence time *τ* of a single Zur protein at a chromosomal tight binding site (Fig. 2a). Each *τ* starts when *r* drops below *r*_0_ and ends when *r* jumps above *r*_0_ (e.g., *τ*’s in Fig. 2a), which are expected to reflect dominantly protein unbinding from DNA, or when the mEos3.2-tag photobleaches/blinks.

**Fig. 2.**
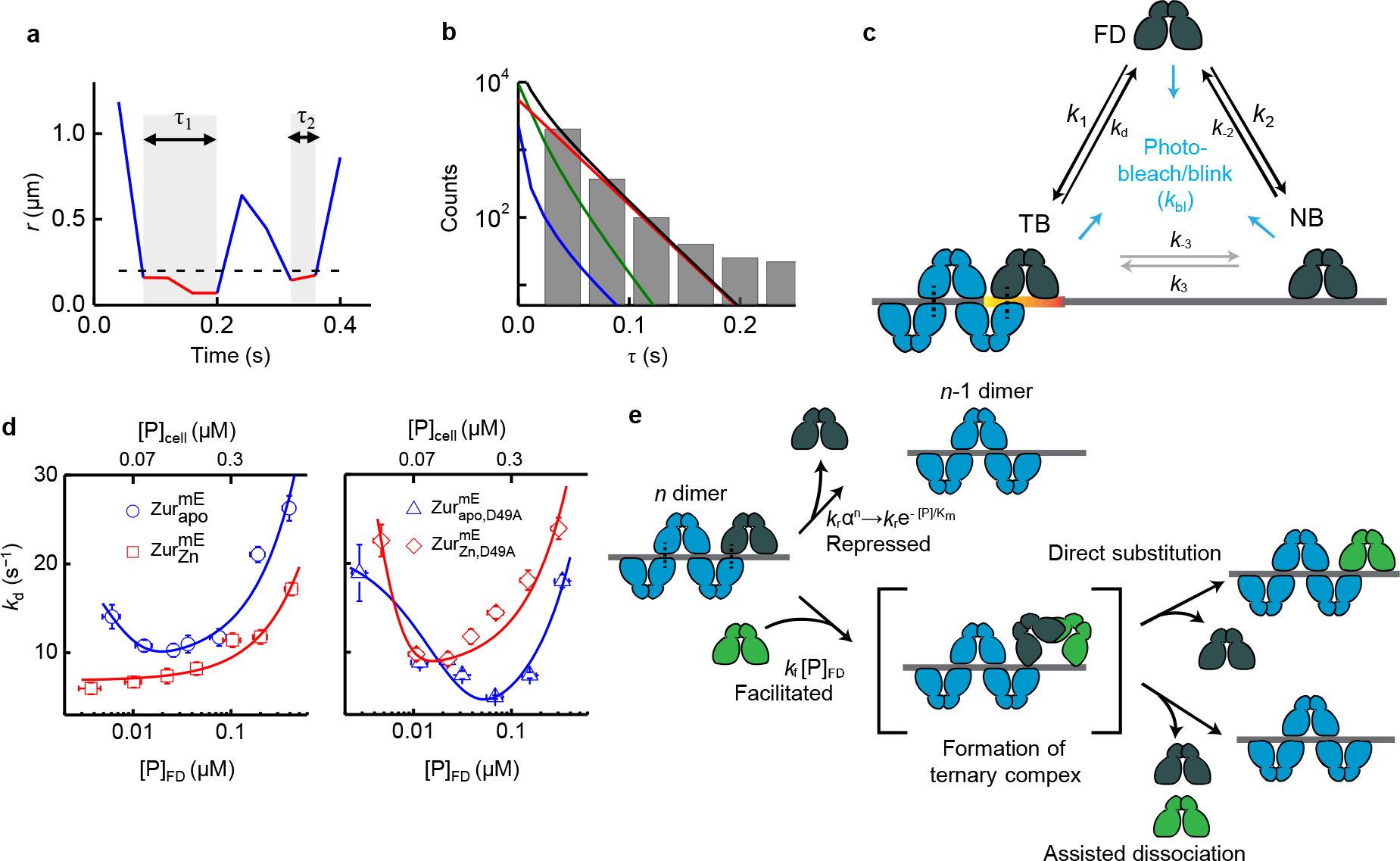
Biphasic unbinding kinetics of Zur from TB sites on chromosome. **a**, Time trajectory of displacement length *r* per time-lapse from a single 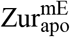 protein. Two microscopic residence time *τ* shown in gray shades; dashed horizontal line: displacement threshold *r*_o_ = 0.2 μm (vertical dashed line in Fig. 1b). **b**, Histogram of *τ* for 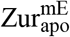 at the cellular concentration of 124 ± 15 nM. Black line: fitting with Eq. (4). Contributions of the three diffusion states are plotted, as color-coded in Fig. 1b-c. **c**, Three-state model of a single Zur protein interacting with DNA in a cell. *k*’s are the rate constants. **d**, Protein-concentration-dependent *k*_d_ for 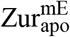 and 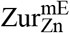 (left) and their corresponding D49A salt-bridge mutants (right). Bottom/top axis refers to free/cellular protein concentration, respectively. Lines are fits with Eq. (2). All error bars are s.d. **e**, Schematic molecular mechanisms for biphasic unbinding of Zur from a TB site. A bound Zur protein (dark blue) within an oligomer on DNA can unbind following either a repressed pathway (top) due to the presence of (*n*-1) proteins nearby or a facilitated pathway (bottom) upon binding another protein (green) to form an intermediate ternary complex, which then proceeds through direct substitution or assist dissociation pathway. Black dashed lines denote salt-bridge interactions.

We analyzed trajectories from many cells of similar cellular Zur concentrations to obtain their corresponding distribution of *τ* (Fig. 2b). We used a quantitative three-state model (i.e., FD, NB, and TB states; Fig. 2c) to analyze the distribution of *τ*, in which the contributions of FD and NB states are deconvoluted (Eq. (4); approximations and validations of this model in Supplementary Note 5)^11^. This model also accounts for mE photobleaching/blinking kinetics, determined from the fluorescence on-time distribution of SMT trajectories (Supplementary Fig. 8). This analysis gave *k*_d_, the apparent first-order unbinding rate constant of Zur from a tight binding site on the chromosome, for each group of cells having similar cellular Zur concentrations.

Strikingly, *k*_d_ for 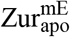 shows a biphasic, repressed-followed-by-facilitated behavior: it initially decreases with increasing free (or total) cellular Zur concentration (i.e., repressed), reaching a minimum at ~130 nM; it then increases toward higher protein concentrations (i.e., facilitated; Fig. 2d, left, blue points). This biphasic behavior is also apparent in the simple averages of residence time 〈*τ*〉 or by analyzing the distributions of *τ* that merely takes into account mE photobleaching/blinking (Supplementary Fig. 9a). The facilitated unbinding of 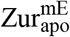 is analogous to those of CueR and ZntR, two MerR-family metalloregulators that we discovered *in vitro* and in living cells^10, 11^; the repressed unbinding of 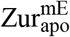 is a *first-of-its-kind* discovery, however.

In contrast, *k*_d_ for 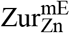 only shows the facilitated unbinding within the accessible cellular protein concentration range (~30 to ~900 nM) — it increases consistently with increasing cellular protein concentrations (Fig. 2d, left, red points). The different behaviors of 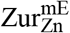 from that of 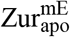 indicate that we could indeed observe the behaviors of the holo-repressor.

### Mechanism of biphasic unbinding of Zur from DNA

Amid the biphasic unbinding of Zur from DNA (Fig. 2d, left), the concentration-facilitated unbinding at higher protein concentrations is analogous to those of CueR and ZntR^11^. There it stems from an assisted dissociation pathway, in which an incoming protein from solution helps an incumbent protein on DNA to unbind, or a direct substitution pathway, in which the incoming protein directly replaces the incumbent one (Fig. 2e, lower)^10, 11^. The rates of both pathways depend linearly on the free protein concentration, and both likely occur through a common ternary protein_2_-DNA complex, in which the two homodimeric proteins each use one DNA-binding domain to bind to half of the dyad recognition sequence^5, 24^. As Zur is also a homodimer, Zur also could form this ternary complex and undergo assisted dissociation or direct substitution, leading to its concentration-facilitated unbinding from DNA.

Regarding the repressed unbinding of apo-Zur in the lower concentration regime, we propose that it likely results from protein oligomerization around the DNA binding site, in which the number of proteins in the oligomer increases with increasing protein concentration and the resulting protein-protein interactions contribute to additional stabilization, thereby repressing protein unbinding rate (Fig. 2e, upper). (The facilitated unbinding later takes over when the protein concentration reaches a high enough level.) Two evidences support our oligomerization proposal: (1) Crystallography showed that two *E. coli* Zur dimers can bind to a short cognate DNA sequence^15^. (2) DNA footprinting showed that *S. coelicoror* Zur forms oligomers around its recognition sites, containing greater than 4 dimers^25^.

To further support this oligomerization proposal, we examined the spatial distribution in the cell of Zur’s residence sites at its TB state; these residence sites correspond to the *r*_0_-thresholded small displacements (Fig. 2a; Supplementary Note 8). For comparison, we further simulated an equal number of sites randomly distributed in a cell of the same size (Supplementary Note 8.1). We then examined their pair-wise distance distributions (PWD), in which Zur oligomerization at chromosomal binding sites should lead to more populations at shorter pair-wise distances. This PWD for 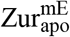 indeed shows a higher population at distances shorter than ~500 nm relative to the simulated random sites (Fig. 3a). However, at the distance scale of a few hundred nanometers, the compaction of chromosome also contributes to the PWD of residence sites^11^. To decouple the contribution of protein oligomerization from chromosome compaction, we examined the fraction of residence sites within a radius threshold *R*. At small *R* (e.g., <100 nm), the contribution of Zur oligomerization to this fraction should dominate over chromosome compaction, as oligomerization is at molecular scale whereas the most compact chromosome in a *E. coli* cell is still around hundreds of nanometer in dimension^11, 26^. At any specified *R* (e.g., 200 nm), the fraction of 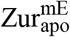 residence sites within the radius *R* increases expectedly with increasing cellular protein concentrations (Fig. 3b, red points), because higher protein concentrations gave higher sampling frequency of residence sites. More important, at lower *R* (e.g., 100 nm), the fraction of 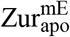 residence sites is larger than that of simulated random sites (Fig 3b, red vs. blue points), and their ratio is larger at lower protein concentrations (Fig. 3b, green points). The average ratio of the fraction of 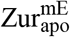 residence sites over that of the simulated random sites is always greater than 1, and it becomes larger at smaller *R* down to <70 nm (Fig. 3c; note our molecular localization precision is ~20 nm; Supplementary Note 3), supporting 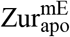 oligomerization at chromosomal tight binding sites at the nanometer scale.

**Fig. 3.**
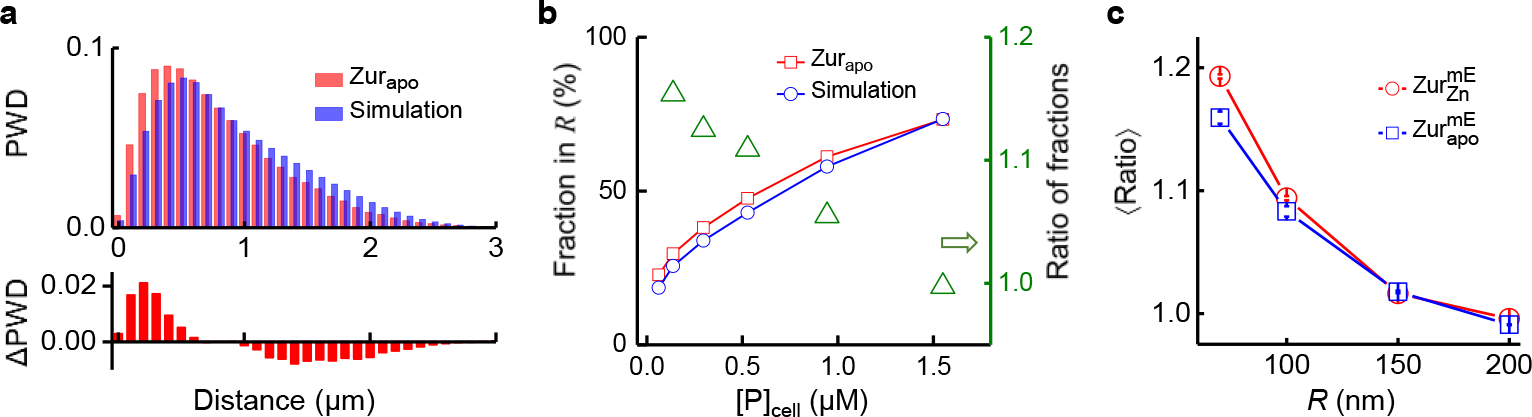
Spatial analysis of Zur’s residence sites. **a**, Normalized pair-wise distance distributions (PWD) of residence sites for 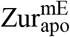 and for simulated random sites in the cell (top), and the difference of 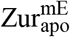 from simulation (bottom). **b**, Fraction of residence sites within a radius threshold *R* (= 100 nm, left axis) for 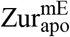 and for simulated random sites as a function of cellular protein concentration. Their ratio (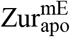 vs. simulation) is plotted against the right axis. **c**, Dependence of the average ratio in **b** across all protein concentrations as a function of the radius threshold *R* for 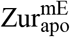 and 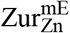.

We formulated a quantitative kinetic model to describe the biphasic unbinding of 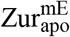. It considers both oligomerization at a TB site and facilitated unbinding via a ternary protein_2_-DNA complex (Fig. 2c and e; Supplementary Note 6). The microscopic unbinding rate constant 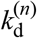 from a TB site with *n* 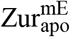 dimers bound as an oligomer comprises three terms:

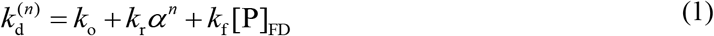

*k*_o_ is a first-order intrinsic unbinding rate constant. The *k*_*r*_*α*^*n*^ term accounts for the repressed unbinding from protein oligomerization, where a first-order rate constant *k*_r_ is attenuated by the factor *α*(0 < *α* < 1) to the exponent of *n*, which depends on the cellular protein concentration and has a maximal value of *n*_0_, the oligomerization number. The third term describes the facilitated unbinding, with *k*_f_ being a second-order rate constant and [P]_FD_ being the concentration of freely diffusing Zur dimers in the cell, as reported for CueR/ ZntR^11^. In the limit of weak oligomerization and low free protein concentrations, the apparent unbinding rate constant *k*_d_ from any TB site is:

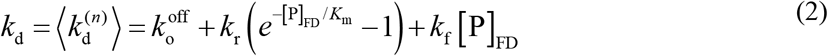

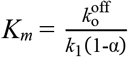; it has the units of protein concentration, reflecting the effective dissociation constant of the protein oligomer on the chromosome. 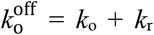; it is a first-order spontaneous unbinding rate constant at the limit of zero cellular protein concentration. Equation (2) satisfactorily fits the biphasic unbinding kinetics of 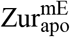 (Fig. 2d, left), giving the associated kinetic parameters (Table 1 and Supplementary Table 6). In particular, *K*_m_ of 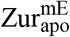 is ~5 nM, indicating that apo-Zur can oligomerize on chromosome at its physiological concentrations in the cells (Fig. 4a).

**Table 1.** Kinetic and thermodynamic parameters for Zur-DNA interaction in *E.coli* cells

|  |  |  |  |  |  |
| --- | --- | --- | --- | --- | --- |
|  | Zur<sup>mE</sup> | Zur<sup>mE</sup><sub>apo</sub> | Zur<sup>mE</sup><sub>Zn</sub> | Zur<sup>mE</sup><sub>apo, D49A</sub> | Zur<sup>mE</sup><sub>Zn, D49A</sub> |
| $k_1$ (nM<sup>-1</sup> s<sup>-1</sup>)<sup>a</sup> | 1.90 ± 0.17 | 1.84 ± 0.20 | 1.10 ± 0.18 | 1.61 ± 0.58 | 1.30 ± 0.19 |
| $k_o^{\text{off}}$ (s<sup>-1</sup>) | 25 ± 12 | 22 ± 21 | 5.4 ± 0.6 | 22.1 ± 1.5 | 36 ± 41 |
| $k_r$ (s<sup>-1</sup>) | 16 ± 11 | 12 ± 20 | n/o<sup>b</sup> | 20.8 ± 1.3 | 27 ± 40 |
| $k_f$ (nM<sup>-1</sup> s<sup>-1</sup>) | 0.028 ± 0.005 | 0.044 ± 0.007 | 0.026 ± 0.033 | 0.049 ± 0.014 | 0.062 ± 0.010 |
| $K_m$ (nM) | 6.0 ± 4.0 | 4.9 ± 7.3 | n/o<sup>b</sup> | 16.2 ± 7.5 | 3.2 ± 1.9 |
| $K_{d1}$ (= $k_o^{\text{off}}/k_1$) (nM)<sup>a</sup> | 12.9 ± 6.2 | 11.7 ± 11.2 | 4.9 ± 1.2 | 13.7 ± 5.0 | 28 ± 20 |
| $K_{d2}$ (= $k_{-2}/k_2$) (nM)<sup>a</sup> | 417 ± 35 | 348 ± 84 | 534 ± 148 | 209 ± 69 | 532 ± 134 |
| $K_{d3}$ (= $k_{-3}/k_3$)<sup>a</sup> | 0.011 ± 0.002 | 0.023 ± 0.007 | 0.022 ± 0.023 | 0.032 ± 0.062 | 0.008 ± 0.006 |
| [D<sub>0</sub>]<sub>NB</sub> (nM)<sup>a</sup> | 1144 ± 84 | 961 ± 205 | 1201 ± 287 | 858 ± 230 | 1538 ± 353 |
| [D<sub>0</sub>]<sub>TB</sub>·$n_o$ (nM)<sup>a</sup> | 42.56 ± 0.94 | 34.3 ± 3.2 | 54 ± 14 | 31.6 ± 5.1 | 38.8 ± 3.8 |

<sup>a</sup> $n_o = 5$ was used in fitting. <sup>b</sup> Not observed

**Fig. 4.**
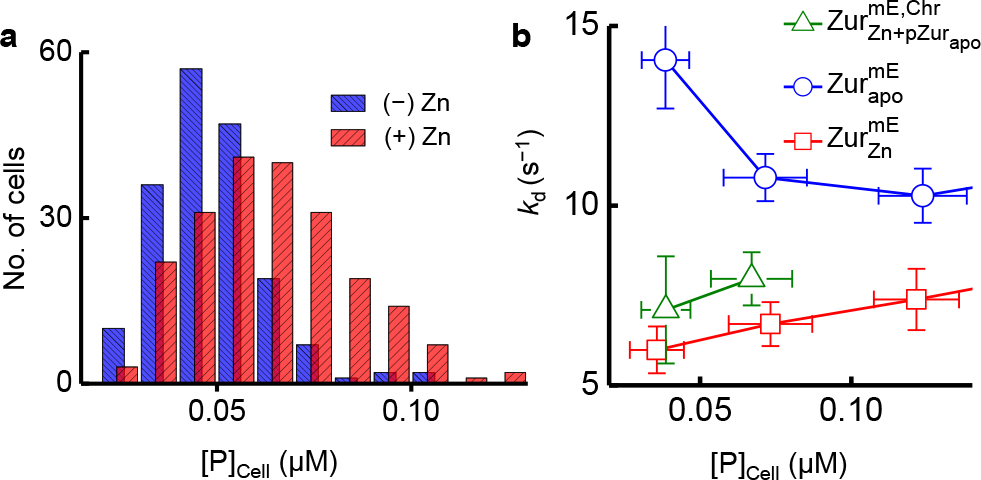
Zur behaviors within the physiological range of cellular protein concentrations. **a**, Distribution of the chromosomally expressed Zur^mE^ concentration in the cell with (+) and without (−) Zn stress in the medium. **b**, Dependence of *k*_d_ on the protein concentration in the cell for 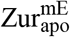, 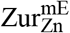, and for 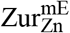 together with a plasmid expressing Zur_apo_ (i.e. 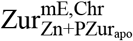) when the mE-tagged Zur is only encoded on the chromosome. The blue circles and red squares for 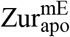 and 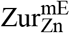 are part of data in Fig. 2d (left).

The same model also allowed for analyzing the relative populations of FD, NB, and TB states of Zur across all cellular protein concentrations, giving additional thermodynamic and kinetic parameters (Table 1, and Supplementary Table 6). Strikingly, the dissociation constant *k*_d1_ of 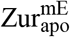 at TB sites of DNA is ~11 nM, merely ~2 times weaker than that of 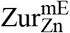 (*k*_d1_ ~5 nM). This is *not* expected because apo-Zur, in both *E. coli* and *B. subtilis*, was shown to have no significant affinity to the consensus sites recognized by holo-Zur^15, 23^. Therefore, the high affinity of 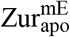 at the TB state suggests that inside cells, apo-Zur likely bind tightly to other, non-consensus sites in the chromosome.

This likelihood is supported by a ChIP-seq analysis in *B. subtilis*, which showed Zur can bind tightly to many locations in the chromosome that do not share consensus with the known recognition sites (although it was undefined whether the detected bindings there were by apo- or holo-Zur)^27^.

### Molecular basis of repressed unbinding

Our model of Zur oligomerization at TB sites was based partly on the structure of two holo-Zur dimers bound to a cognate DNA, which showed two inter-dimer D49-R52 salt bridges^15^. To probe the role of these salt bridges in Zur oligomerization, we made the D49A mutation, known to disrupt the interactions^15^. For apo-Zur, the resulting mutant 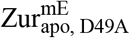 still exhibits the biphasic unbinding behavior, however the minimum of the apparent unbinding rate constant *<u>k</u>*_d_ shifted to a higher cellular protein concentration (Fig. 2d, right). Its *K*_m_ is 16.2 ± 7.5 nM, three times larger than that of 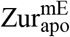 (Table 1), indicating a weakened oligomerization affinity and thus a significant role of these salt bridges.

More strikingly, for 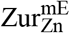, which only showed facilitated unbinding (Fig. 2d, left), the resulting mutant 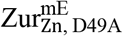 clearly shows biphasic unbinding with *K*_m_ = 3.2 ± 1.9 nM (Fig. 2d, right; Table 1). Therefore, holo-Zur also possesses repressed unbinding kinetics — it was invisible for 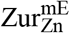 likely because its *K*_m_ is smaller than the low limit of accessible cellular protein concentrations (~3 nM), but emerges after the D49A mutation, which further supports the importance of the salt bridges in Zur oligomerization and repressed unbinding behaviors.

## DISCUSSION

We have uncovered that the Fur-family Zn^2+^-sensing transcription regulator Zur exhibits two unusual behaviors that challenge conventional paradigms of regulator-chromosome interactions. First, apo-Zur, the non-repressor form and a long-presumed non-DNA binder, can actually bind to chromosome tightly, likely at different locations from the consensus sequence recognized by holo-Zur, the repressor form. This tight chromosome binding by apo-Zur challenges the paradigm of regulator on-off model for transcription repression (or activation)^1, 2^. Second, the unbinding kinetics of both apo- and holo-Zur not only exhibit facilitated unbinding, a newly discovered phenomenon for a few DNA-binding proteins^6, 7, 9, 28^, but also show repressed unbinding, a *first-of-its-kind* phenomenon that likely results from Zur oligomerization on chromosome, facilitated by inter-dimer salt bridges. Overall, Zur has biphasic unbinding kinetics from chromosome with increasing cellular protein concentrations, which challenges the paradigm of protein unbinding being typically unimolecular processes whose first-order rate constants do not depend on the protein concentration.

To probe whether the biphasic unbinding of Zur occurs within the physiological cellular protein concentrations, we quantified cellular Zur^mE^ concentration when it is encoded only at the chromosomal locus (Fig. 4a). In minimal medium without Zn stress, the cellular Zur^mE^, which is mostly in the apo-form, ranges from ~24 to 108 nM (mean = 50 ± 14 nM), within which apo-Zur unbinding from TB sites is in the repressed unbinding regime and slows down by ~42% from the lowest to the highest protein concentration (Fig. 4b). When stressed by 20 μM Zn^2+^, the cellular Zur^mE^, now mostly in the holo-form, ranges from ~26 to 124 nM (mean = 63 ± 20 nM), reflecting an average of ~28% protein concentration increase induced by Zn stress. In this protein concentration range, holo-Zur is already in the facilitated unbinding regime, and its unbinding rate from a recognition site can increase by ~36% (Fig. 4b).

Within the physiological protein concentration range, the opposite dependences of unbinding kinetics on the cellular protein concentration between apo- and holo-Zur could provide functional advantages for an *E. coli* cell to repress or de-repress Zn uptake genes. When cell encounters environmental Zn stress that demands strong repression of Zn uptake, the cellular concentration of Zur swings upward and it becomes dominantly in the holo-repressor form. The unbinding of holo-repressor from recognition sites could be facilitated by its increasing concentration (Fig. 5a), but the facilitated unbinding via direct substitution by another holo-repressor has no functional consequence while facilitated unbinding via assisted dissociation will be immediately compensated by a rebinding of a holo-repressor (the rebinding would occur within ~0.014 s; Supplementary Note 7). For those cellular Zur in the apo non-repressor form, its unbinding from DNA slows down, keeping them longer (i.e., stored) at non-consensus chromosomal sites (Fig. 5b). On the other hand, when cell transitions to a Zn-deficient environment that demands derepression of Zn uptake, the cellular Zur protein concentration goes down. Here unbinding of the holo-repressor would be slower (Fig. 5c), which is undesirable for derepression, while the unbinding of the apo-form would become faster, releasing them from the non-consensus “storage” sites on the chromosome into the cytosol (Fig. 5d). If the cytosolic apo-Zur could possibly facilitate the unbinding of holo-Zur from promoter recognition sites (e.g., through assisted dissociation), it would give a more facile transition to derepression. To support this possibility, we measured the apparent unbinding rate constant *k*_d_ for chromosomally encoded 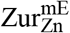 in cells that contains a plasmid encoding an untagged Zur_apo_ mutant (i.e., C88S). When the expression of this Zur_apo_ mutant is induced, *k*_d_ of 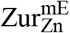 increases by ~28% at any cellular 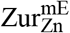 concentration (Fig. 4b, green vs. red points), indicating that apo-Zur can indeed facilitate the unbinding of holo-Zur from recognition sites (Fig. 5e).

**Fig. 5.**
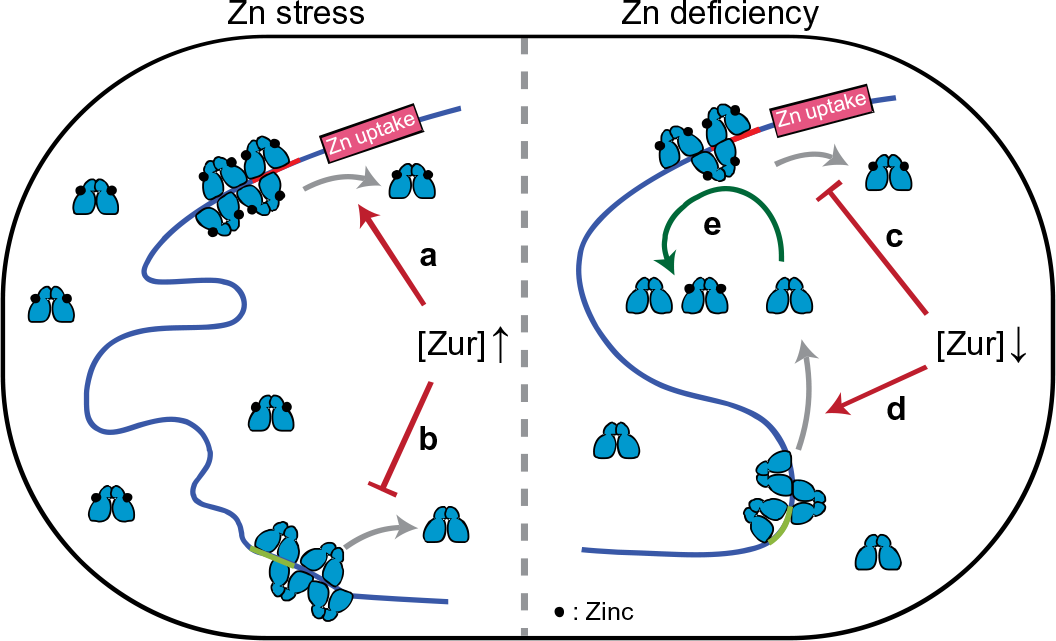
Functional model of holo- and apo-Zur unbinding behaviors in *E.coli* upon encountering zinc stress or deficiency. Upon zinc stress, unbinding of holo-Zur from operator site is facilitated (**a**) while that of apo-Zur from storage site is repressed (**b**) due to increase in cellular protein concentration. Upon zinc deficiency, the facilitated unbinding of holo-Zur is attenuated (**c**) while the unbinding of apo-Zur is less repressed (**d**) due to decrease in cellular protein concentration. Released apo-Zur into cytosol could facilitate holo-Zur to unbind (**e**), helping transition to de-repression of zinc uptake.

Multivalent contacts with DNA, which underlie the facilitated unbinding, and salt-bridge interactions between proteins, which underlie Zur oligomerization and its repressed unbinding, are both common for protein-DNA and protein-protein interactions, respectively^5, 7, 10, 28–36^. Therefore, the biphasic unbinding behavior from DNA discovered here for Zur could be broadly relevant to many other proteins in gene regulation.

## METHODS

### Bacterial strains and sample preparation

All strains were derived from the *E.coli* BW25113 strain as detailed in Supplementary Note 1. Zur^mE^ was either encoded at its chromosomal locus via lambda-red homologous recombination^37^ or in a pBAD24 plasmid in a Δ*zur* deletion strain^38^. Mutant forms of Zur (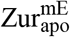, 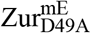, or 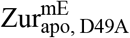) were generated via site-directed mutagenesis in pBAD24, which was introduced into the Δ*zur* strain.

All cell imaging experiments were done at room temperature in M9 medium supplemented with amino acids, vitamins, and 0.4% glycerol. 20 μM ZnSO_4_ was used for Zn stress conditions. The cells were immobilized on an agarose pad in a sample chamber. Details in Supplementary Note 3.

### SMT and SCQPC

SMT and SCQPC were performed on an inverted fluorescence microscope, as reported^11^ (Supplementary Note 3). For SMT, inclined epi-illuminated 405 nm and 561 nm lasers photoconverted and excited single mEos3.2 molecules, respectively. 561 nm excitation-imaging were in stroboscopic mode, with 4 ms laser excitation pulses separated by 40 ms time lapse, synchronized with the camera exposure, so that the mobile proteins still appear as diffraction-limited spots. A custom-written MATLAB software was used to identify diffraction-limited fluorescence spots and fit them with two-dimensional Gaussian functions, giving ~20 nm localization precision^11, 39^. Time trajectories of positions and displacement length *r* between adjacent images were then extracted.

SCQPC was performed after SMT. The remaining proteins were firstly photoconverted to the red form by a long 405 nm laser illumination. The total cell red fluorescence was then imaged by the 561 nm laser to determine the protein copy number, provided the average fluorescence of a single mEos3.2 from the earlier SMT. The photoconversion efficiency of mEos3.2^40^ and dimeric state of Zur were accounted for. Cell volumes were determined by fitting their optical transmission image contours with the model geometry of a cylinder with two hemispherical caps.

### Resolution of diffusion states

The effective diffusion constants and the fractional populations of diffusion states were extracted by analyzing the CDF of displacement length *r* per time-lapse (*T*_tl_ = 40 ms), using a linear combination of three diffusion terms of CDF, as reported^11^ (Equation (3)). Each term is from a 2-D Brownian diffusion model^18, 41, 42^, which was regularly used to analyze SMT results of proteins in bacterial and mammalian cells^18, 21, 42–46^ (model justification in Supplementary Note 4).

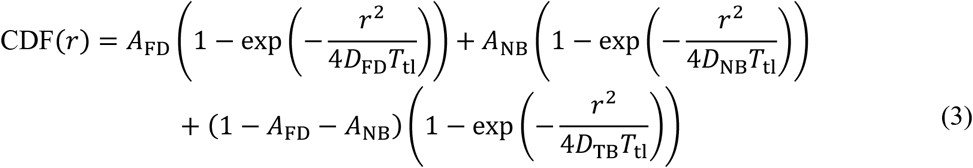

We globally fitted the CDFs across groups of cells of different cellular protein concentrations, in which the diffusion constants (*D*’s) of respective diffusion states were shared but their factional populations (*A*’s) were allowed to vary. Three terms were always the minimal number of diffusion states to satisfactorily fit the CDF (details in Supplementary Note 4 and Supplementary Tables 4-5).

Note these diffusion constant values are not the intrinsic ones, as they are influenced by the cell confinement effect^47^, which decreases the magnitude of the apparent diffusion constant, and by the time-lapse effect of imaging, where longer time lapse gives apparently smaller diffusion constants; both of these effects are most significant on the FD state, less on the NB state, and negligible on the TB state, and were evaluated quantitatively in a previous study of metal-responsive transcription regulators of a different family^11^.

### Determination and analysis of *k*_d_

A three-state (FD, NB, and TB state) kinetic model, including the interconversion between states and photobleaching/blinking rates (Fig. 2c), was used to analyze the distribution of residence times (upper thresholded by *r*_0_; Fig. 2a) at chromosomal TB sites to extract the apparent unbinding rate constant *k*_d_. The respective residence time distribution functions *φ*(*τ*) for the FD, NB, and TB states with given diffusion constants (*D*’s), the unbinding rate constant from the NB state *k*_−2_, and photobleaching/blinking rate constant *k*_bl_ were derived to fit the *τ* distribution with the overall distribution function *φ*_all_(*τ*) (Eq. (4); Supplementary Note 5).

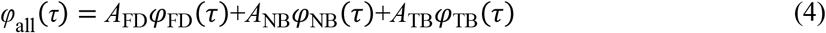

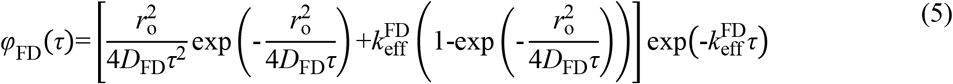

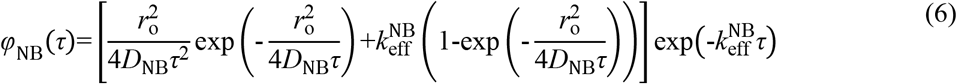

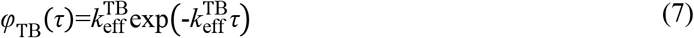

Here 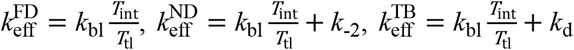, and *A_i_* is the fractional population of *i*^th^-state.

The dependence of *k*_d_ on the cellular free diffusing protein concentration [P]_FD_ was analyzed with Eq. (2), containing three terms representing spontaneous, repressed, and facilitated unbinding with the corresponding rate constants 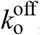, *k*_r_, and *k*_f_, respectively (derivation in Supplementary Note 6).

### Analysis of relative populations

The same three-state kinetic model (Fig. 2c) was used to analyze the relative populations of FD, NB, and TB states of Zur across all cellular protein concentrations. Oligomerization/deoligomerization of Zur at a TB site was modeled as 1-D sequential binding/unbinding, analogous to the Brunauer-Emmett-Teller multilayer-adsorption theory^48^ but with a limited number *n*_0_ of binding site and merely one binding rate constant *k*_1_ (see Supplementary Note 7 for detailed derivation). Quasi-equilibrium approximation of interconversion among states was used, which approximates that the timescale of interconversion between states (~ms) are much shorter than the experimental imaging time (~hours). The kinetic parameters are then related to the relative concentrations of the proteins at three diffusion states.

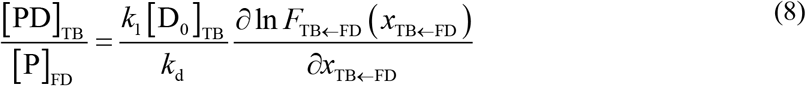

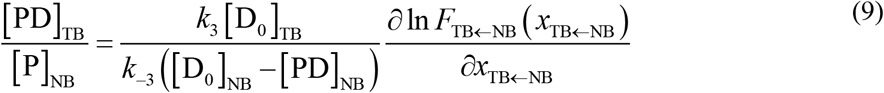

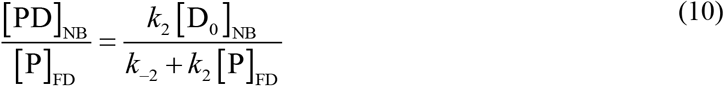

Here [P]_FD_, [PD]_NB_, and [PD]_TB_ are the cellular protein concentrations of FD, NB, and TB states, respectively. 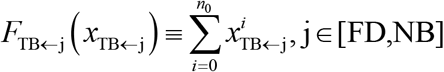, where 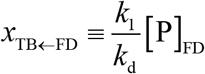 and 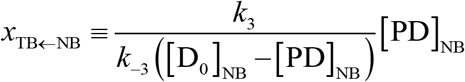. [D_o_]_TB_ and [D_o_]_NB_ are the effective cellular concentrations of TB and NB sites, respectively. Thermodynamic quantities such as the dissociation constant of TB 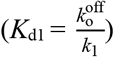 and 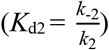 were also determined from this analysis.

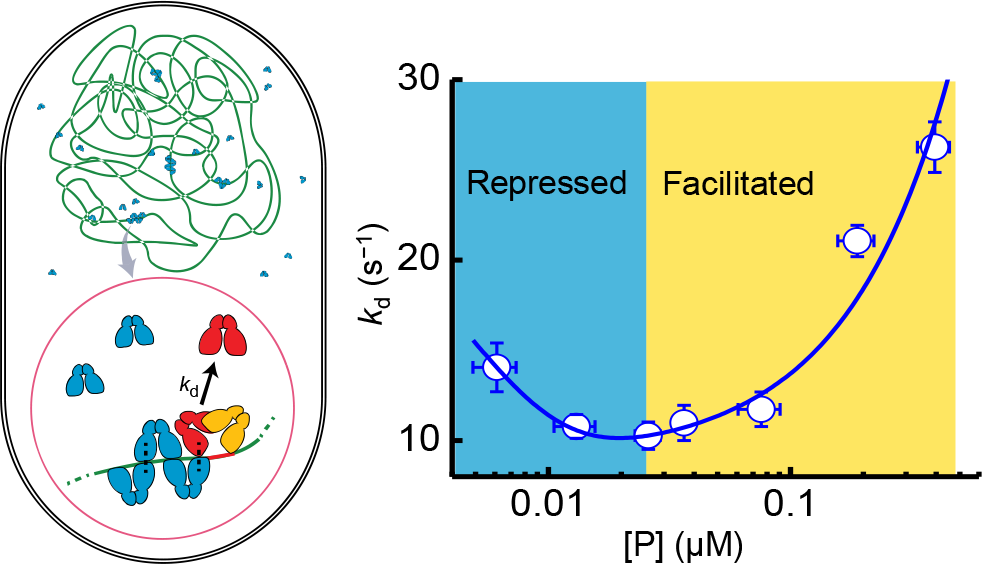
TOC Graph.

## Acknowledgements

This research is supported by the National Institutes of Health (GM109993). We thank A. G. Santiago for providing molecular biology protocols and materials, T.-Y. Chen for providing iqPALM MATLAB codes, Y. Aye and C. Kinsland for access to biology facilities, and J. D. Helmann for discussion.

## Author contributions

W.J. and P.C. designed research; W.J. performed experiments, derived theory, coded software, and analyzed data; W.J. and P.C. discussed the results and wrote the manuscript.

## Competing interests

The authors declare no competing interest.

## Additional information

**Supplementary information** is available for this paper.

## REFERENCES

1. Schramm, L. & Hernandez, N. Recruitment of RNA polymerase III to its target promoters. Genes Dev. 16, 2593–2620 (2002).

2. Deuschle, U., Gentz, R. & Bujard, H. lac Repressor blocks transcribing RNA polymerase and terminates transcription. Proc. Natl Acad. Sci. USA 83, 4134–4137 (1986).

3. Rajagopalan, S. et al. Studies of IscR reveal a unique mechanism for metal-dependent regulation of DNA binding specificity. Nat. Struct. Mol. Biol. 20, 740–747 (2013).

4. Nesbit, A. D., Giel, J. L., Rose, J. C. & Kiley, P. J. Sequence-specific binding to a subset of IscR-regulated promoters does not require IscR Fe-S cluster ligation. J. Mol. Biol. 387, 28–41 (2009).

5. Chen, T. Y., Cheng, Y. S., Huang, P. S. & Chen, P. Facilitated Unbinding via Multivalency-Enabled Ternary Complexes: New Paradigm for Protein-DNA Interactions. Acc. Chem. Res. 51, 860–868 (2018).

6. Graham, J. S., Johnson, R. C. & Marko, J. F. Concentration-dependent exchange accelerates turnover of proteins bound to double-stranded DNA. Nucleic Acids Res. 39, 2249–2259 (2011).

7. Gibb, B. et al. Concentration-dependent exchange of replication protein A on single-stranded DNA revealed by single-molecule imaging. PloS One 9, e87922 (2014).

8. Geertsema, H. J., Kulczyk, A. W., Richardson, C. C. & van Oijen, A. M. Single-molecule studies of polymerase dynamics and stoichiometry at the bacteriophage T7 replication machinery. Proc. Natl Acad. Sci. USA 111, 4073–4078 (2014).

9. Lewis, J. S. et al. Single-molecule visualization of fast polymerase turnover in the bacterial replisome. eLife 6 (2017).

10. Joshi, C. P. et al. Direct substitution and assisted dissociation pathways for turning off transcription by a MerR-family metalloregulator. Proc. Natl Acad. Sci. USA 109, 15121–15126 (2012).

11. Chen, T. Y. et al. Concentration- and chromosome-organization-dependent regulator unbinding from DNA for transcription regulation in living cells. Nat. Commun. 6, 7445 (2015).

12. Hantke, K. Bacterial zinc uptake and regulators. Curr. Opin. Microbiol. 8, 196–202 (2005).

13. Hemm, M. R. et al. Small stress response proteins in Escherichia coli: proteins missed by classical proteomic studies. J. Bacteriol. 192, 46–58 (2010).

14. Panina, E. M., Mironov, A. A. & Gelfand, M. S. Comparative genomics of bacterial zinc regulons: enhanced ion transport, pathogenesis, and rearrangement of ribosomal proteins. Proc. Natl Acad. Sci. USA 100, 9912–9917 (2003).

15. Gilston, B. A. et al. Structural and mechanistic basis of zinc regulation across the E. coli Zur regulon. PLoS Biol. 12, e1001987 (2014).

16. Zhang, M. et al. Rational design of true monomeric and bright photoactivatable fluorescent proteins. Nat. Methods 9, 727–729 (2012).

17. McKinney, S. A. et al. A bright and photostable photoconvertible fluorescent protein. Nat. Methods 6, 131–133 (2009).

18. Elf, J., Li, G. W. & Xie, X. S. Probing transcription factor dynamics at the single-molecule level in a living cell. Science 316, 1191–1194 (2007).

19. Javer, A. et al. Short-time movement of E. coli chromosomal loci depends on coordinate and subcellular localization. Nat. Commun. 4, 3003 (2013).

20. Mehta, P. et al. Dynamics and stoichiometry of a regulated enhancer-binding protein in live Escherichia coli cells. Nat. Commun. 4, 1997 (2013).

21. Uphoff, S. et al. Single-molecule DNA repair in live bacteria. Proc. Natl Acad. Sci. USA 110, 8063–8068 (2013).

22. Mazza, D., Ganguly, S. & McNally, J. G. Monitoring dynamic binding of chromatin proteins in vivo by single-molecule tracking. Methods Mol. Biol. 1042, 117–137 (2013).

23. Ma, Z., Gabriel, S. E. & Helmann, J. D. Sequential binding and sensing of Zn(II) by Bacillus subtilis Zur. Nucleic Acids Res. 39, 9130–9138 (2011).

24. Chen, P. et al. Single-molecule dynamics and mechanisms of metalloregulators and metallochaperones. Biochemistry 52, 7170–7183 (2013).

25. Choi, S. H. et al. Zinc-dependent regulation of zinc import and export genes by Zur. Nat. Commun. 8, 15812 (2017).

26. Wang, W. et al. Chromosome organization by a nucleoid-associated protein in live bacteria. Science 333, 1445–1449 (2011).

27. Prestel, E., Noirot, P. & Auger, S. Genome-wide identification of Bacillus subtilis Zur-binding sites associated with a Zur box expands its known regulatory network. BMC Microbiol. 15, 13 (2015).

28. Hadizadeh, N., Johnson, R. C. & Marko, J. F. Facilitated Dissociation of a Nucleoid Protein from the Bacterial Chromosome. J. Bacteriol. 198, 1735–1742 (2016).

29. Geertsema, H. J., Kulczyk, A. W., Richardson, C. C. & van Oijen, A. M. Single-molecule studies of polymerase dynamics and stoichiometry at the bacteriophage T7 replication machinery. Proc. Natl Acad. Sci. USA 111, 4073–4078 (2014).

30. Norel, R., Sheinerman, F., Petrey, D. & Honig, B. Electrostatic contributions to protein-protein interactions: fast energetic filters for docking and their physical basis. Protein Sci. 10, 2147–2161 (2001).

31. Xu, D., Tsai, C. J. & Nussinov, R. Hydrogen bonds and salt bridges across protein-protein interfaces. Protein Eng. 10, 999–1012 (1997).

32. Zhang, Z., Witham, S. & Alexov, E. On the role of electrostatics in protein-protein interactions. Phys. Biol. 8, 035001 (2011).

33. Persson, B. A. & Lund, M. Association and electrostatic steering of alpha-lactalbumin-lysozyme heterodimers. Phys. Chem. Chem. Phys. 11, 8879–8885 (2009).

34. Gunasekaran, K. et al. Enhancing antibody Fc heterodimer formation through electrostatic steering effects: applications to bispecific molecules and monovalent IgG. J. Biol. Chem. 285, 19637–19646 (2010).

35. Persson, B. A., Jonsson, B. & Lund, M. Enhanced protein steering: cooperative electrostatic and van der Waals forces in antigen-antibody complexes. J. Phys. Chem. B 113, 10459–10464 (2009).

36. Hemsath, L. et al. An electrostatic steering mechanism of Cdc42 recognition by Wiskott-Aldrich syndrome proteins. Mol. Cell 20, 313–324 (2005).

37. Datsenko, K. A. & Wanner, B. L. One-step inactivation of chromosomal genes in Escherichia coli K-12 using PCR products. Proc. Natl Acad. Sci. USA 97, 6640–6645 (2000).

38. Guzman, L. M., Belin, D., Carson, M. J. & Beckwith, J. Tight regulation, modulation, and high-level expression by vectors containing the arabinose PBAD promoter. J. Bacteriol. 177, 4121–4130 (1995).

39. Thompson, R. E., Larson, D. R. & Webb, W. W. Precise Nanometer Localization Analysis for Individual Fluorescent Probes. Biophys. J. 82, 2775–2783 (2002).

40. Durisic, N. et al. Single-molecule evaluation of fluorescent protein photoactivation efficiency using an in vivo nanotemplate. Nat. Methods 11, 156–162 (2014).

41. Fick, A. V. On liquid diffusion. The London, Edinburgh, and Dublin Philosophical Magazine and Journal of Science 10, 30–39 (1855).

42. Gebhardt, J. C. et al. Single-molecule imaging of transcription factor binding to DNA in live mammalian cells. Nat. Methods 10, 421–426 (2013).

43. English, B. P. et al. Single-molecule investigations of the stringent response machinery in living bacterial cells. Proc. Natl Acad. Sci. USA 108, E365–373 (2011).

44. Mazza, D. et al. A benchmark for chromatin binding measurements in live cells. Nucleic Acids Res. 40, e119 (2012).

45. Niu, L. & Yu, J. Investigating intracellular dynamics of FtsZ cytoskeleton with photoactivation single-molecule tracking. Biophys. J. 95, 2009–2016 (2008).

46. Mueller, F., Stasevich, T. J., Mazza, D. & McNally, J. G. Quantifying transcription factor kinetics: at work or at play? Crit. Rev. Biochem. Mol. Biol. 48, 492–514 (2013).

47. Chen, T. Y. et al. Quantifying Multistate Cytoplasmic Molecular Diffusion in Bacterial Cells via Inverse Transform of Confined Displacement Distribution. J. Phys. Chem. B 119, 14451–14459 (2015).

48. Brunauer, S., Emmett, P. H. & Teller, E. Adsorption of Gases in Multimolecular Layers. J. Am. Chem. Soc. 60, 309–319 (1938).

